# Rational nanotoolbox with theranostic potential for medicated pro-regenerative corneal implants

**DOI:** 10.1101/581090

**Authors:** Hirak K. Patra, Mohammad Azharuddin, Mohammad M. Islam, Georgia Papapavlou, Suryyani Deb, Geyunjian Harry Zhu, Thobias Romu, Ashis K. Dhara, Mohammad J. Jafari, Amineh Gadheri, Jorma Hinkula, Madhavan S Rajan, Nigel Slater

## Abstract

Cornea diseases are a leading cause of blindness and the disease burden is exacerbated by the increasing shortage around the world for cadaveric donor corneas. Despite the advances in the field of regenerative medicine, successful transplantation of laboratory made artificial corneas has not been fully realised in clinical practice. The causes of failure of such artificial corneal implants are multifactorial and include latent infections from viruses and other micorbes, enzyme over-expression, implant degradation, extrusion or delayed epithelial regeneration. Therefore, there is an urgent unmet need for developing customized corneal implants to suit the host environment, counter the effects of inflammation or infection and that are able to track early signs of implant failure *in situ*. In the present work, we describe a nano toolbox comprising tools for drug release and in addition capable of being infection responsive, promoting regeneration including non-invasive monitoring of *in situ* corneal environment. These nano constructs can be incorporated within pro-regenerative biosynthetic implants, transforming them into theranostic devices able to respond to biological changes following implantation.

## Introduction

Transplantation of solid organs suffers from a dire shortage of donated tissue and organs, and issues of post-operative infection and immune rejection leading to graft failure. In corneal transplantation, the most common transplantation procedure, there is a shortfall of corneas that leaves an estimated 69 out of 70 patients with treatment delay and a waiting list of 12.7 million patient worldwide^1^. This does not include patients in remote areas, or those that may not be prioritized for grafting due to high risk of rejection. Once a patient rejects a donor graft, the chances of the next graft being rejected are much higher, so the probability of each successive graft failing increases resulting in patients moving closer towards irreversible blindness^2^. Cornea transplantation is the only widely accepted treatment for restoring eyesight. However, corneas with severe pathologies, e.g. those causing inflammation or keratitis and severe vascularisation, are at high risk for rejecting conventional human donor transplantation^3^.

The most common infectious cause for blindness worldwide is Herpes Simplex Keratitis (HSK), which is caused by the herpes simplex virus serotype 1 (HSV-1). In the USA alone, almost 500,000 cases of corneal HSV infection were reported with new and recurrent cases each year^4,5^. Approximately 20% of HSV-infected patients will develop HSK where the resulting scarring compromises vision sufficiently to require transplantation to restore eyesight. Patients with HSK are often prescribed lifelong prophylactic drugs to prevent reactivation, as reactivation can lead to graft rejection and failure. Systemic prophylactic drugs when required are often at higher doses to counter the pharmacokinetics of poor absorption through corneal layers and sub therapeutic levels reached within the eye^6,7^. Further, long term prophylaxis is associated with significant side effects and cases of ACV-resistance have been reported^7,8^.

Artificial corneas in the form of prostheses known as keratoprostheses (or KPros) were developed for managing high-risk corneas^9^. One of the most successful KPros is the Boston KPro, comprising a synthetic poly methyl methacrylate (PMMA) optic and front plate. Together with a PMMA/titanium back plate^10^, the front and back plates are used to sandwich a skirt comprising a donor corneal graft. However, even in the Boston KPro, lifelong antibiotics are necessary to prevent infection due to incomplete bio integration of the device^11^ and the device exposes the patients to a very high risk (60-76%) of glaucoma leading to non-reversible blindness. KPros are therefore only used in the case of end-stage disease^12^ and a shunt is often co-implanted to manage glaucoma^13^. These implants are not medicated not do they use materials that are alligned to human corneal collagen. Interestingly, there has been proposed co-implantation of an intraocular pressure sensor embedded to detect glaucoma (ClinicalTrials.org identifier: NCT 03421548) but the trial was withdrawn.

Members of our team recently reported the successful 24 month grafting of cell-free, recombinant human collagen type III-methacryloyloxyethyl phosphorylcholinephospholipid (RHCIII-MPC) implants into keratitis patients^14^. One patient had HSK and the other had fungal keratitis. Both showed marked vision improvement at an average of 24 months post-grafting. Although the HSK patient was doing well at the 2-year check-up, HSV-1 infections can recur at any time due to prolonged viral latency in the trigeminal nerves that supply the cornea ^15^. The surgical procedure and post-operative steroids have additionally been implicated in viral reactivation^16^. A complete theranostic implant system with *in situ* monitoring and detection system for viral reactivation and ability to release drugs to block viral activity is therefore merited.

Our team previously used layer-by-layer nano-coating technology to engineer a contact lens that could detect elevated levels of interleukin-1, one of the earliest cytokines that initiate the inflammatory cascade in HSK^17^. The diagnostic component consisted of an antibody-chromophore colorimetric change in the presence of increased interleukin-1. However, the therapeutic aspect of this theranostic implant system was not well-developed. Therefore, we focus here on developing a complete integrative theranostic system with therapeutics and diagnostic ability. We examined the potential of a therapeutic, pro-regeneration corneal implant that incorporated acyclovir, a widely-used antiviral drug with gold nanoparticles (GNPs) to design a medicated device capable of regulated drug release following transplantation. We have developed and examined the possibility of tracking the drug profile within the implants using minimally invasive magnetic resonance imaging (MRI).We have simultaneously developed another nano tool by replacing the gold particles which might not be readily traced under relatively dry eye conditions, with iron nanoparticles (FeNP) that have been widely used for MRI tracing. We have developed a Nano toolbox that incorporate the theranostic approach to combat major corneal issue by offering tools those are cornea compatible, releasing the appropriate drugs, promoting the growth and to monitor the regeneration process non-invasively to provide patients’ specific solution. We show that these tools are easily incorporated into pro-regeneration corneal constructs, converting these customised artificial cornea into therapeutic implants (Tx). These Tx implants together with previously described diagnostic contact lenses constitute a theranostic system that can be further developed for a personalized clinical application.

## Results

### Design, synthesis and characterization of the theranostic gold nanosystems

#### Acyclovir-conjugated gold nanoparticles (GNPs)

A range of gold nanoparticles (GNPs) were synthesized and one with 20nm hydrodynamic diameter and surface plasmon absorbance maximum at 520nm (Fig. 1a) and surface zeta potential of −54mV were optimized^18^ for our theranostic purpose for ACV conjugation and function. We optimized a non-covalent ionic coupling approach with the zwitterionic forms of ACV (with both the weak acid and basic moieties with pKa= 9.3 for ionization of the guanine OH, and pKb=2.3 for protonation of guanine nitrogen). The Hyperquad Simulation and Speciation (HySS) was used to analyse zwitterion features of ACV to obtain the optimal conditions with the required flexibility for conjugation (Fig. 1b) ^19,20^. Synthesized GNPs surface was capped with polybasic citrate (Fig. S1), which yields hydrogen at pH 6.4 and pH 4.7.

**Figure 1.**
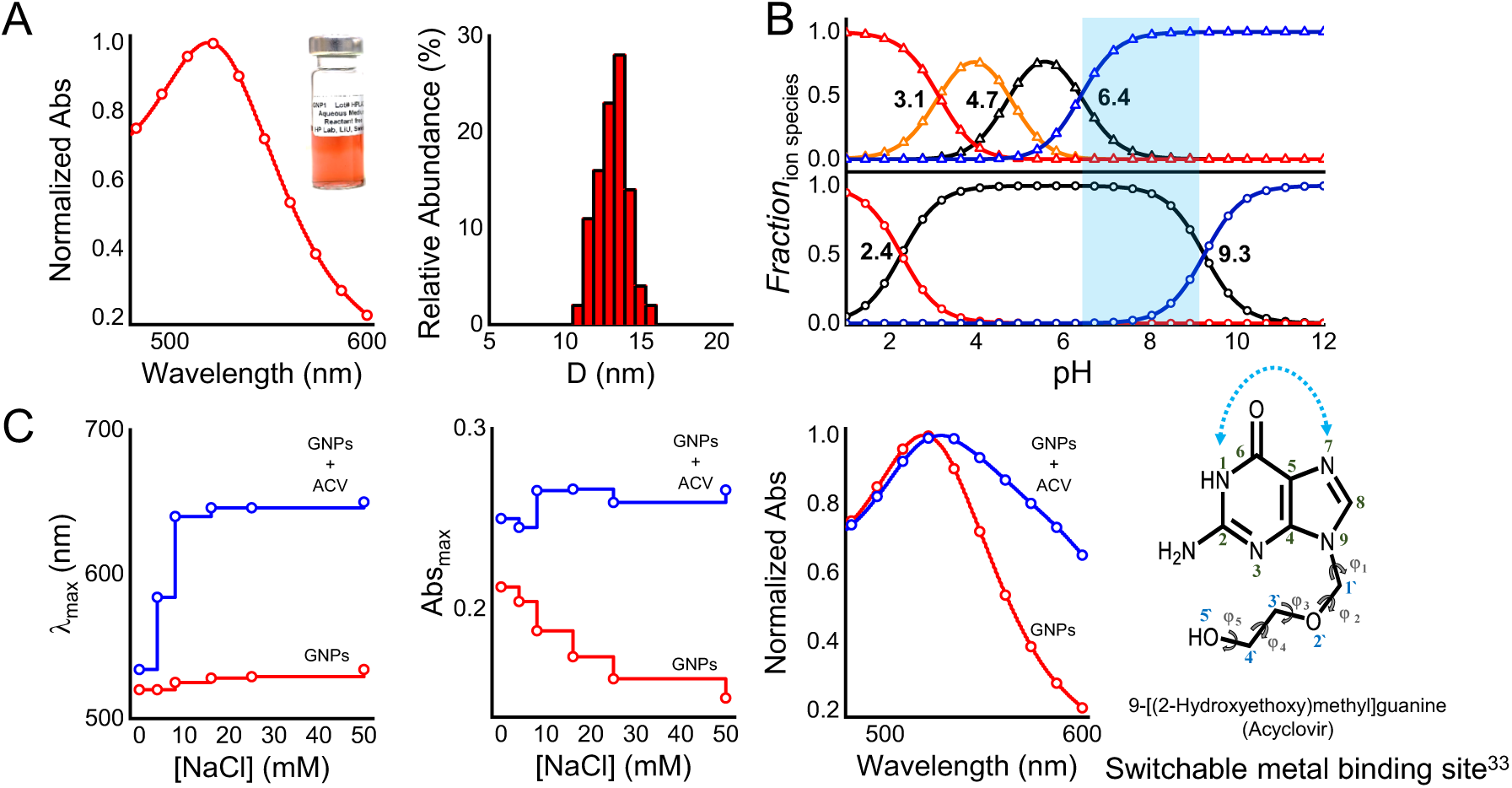
Development of GNP nanosystem. **A**, Left: Absorption spectrum for citrate capped GNPs used in this study. Right: Hydrodynamic diameter (Size) histogram for colloidal GNP solution used (inset). **B**, Top: Hyperquad simulation and speciation (HySS) of citrate and anti-viral drug acyclovir with indicated optimised zone for drug loading with loosely bound semi ionic Lewis bonds^21^. Bottom: The structural flexibility of acyclovir in facilitating conjugation. **C**, Functionalisation of acyclovir on the activated and optimized gold nano-surface by flocculation test (left-middle) and shift in surface Plasmon absorption maxima λ_max_(right).

Therefore, above this pH, the GNP surface becomes negative, enabling ionic interactions with zwitterionic ACV^22^. A pH range from 6.4 to 9 was identified as the zwitterion zone (cyan; Fig. 1b) that facilitated interaction with the activated GNPs surface for ACV loading. Upon conjugation with ACV, the size distribution of GNPs showed shift from 20 nm to ∼81nm in hydrodynamic diameter, with an absorbance maxima at 528nm and zeta potential of −16.6mV. Direct monitoring and conjugation of GNPs with the ACV was studied *in vitro* using a flocculation test^23^ by increasing strength in presence of salt, NaCl (Fig. 1c), showed disruption of the ionic stability^21^ between GNPs and ACV leading to the formation of large particle aggregates as represented by higher red shift in the λ_max_ of the conjugated species as opposed to the unconjugated GNPs (see *Supplementary section 3 for more molecular signature with FTIR*). A high concentration of NaCl perturbs the electrical double layer of the GNPs and induces a shift in the equilibrium between electrostatic repulsion and attraction of the particles^23^.

### Composite collagen Tx implants

Collagen implants comprising 10% (wt/wt) were fabricated as previously described^24^. A detailed description for the fabrication of therapeutic implants (Tx) with ACV-conjugated GNPs (ACV-GNPs) is illustrated in Figure S2. The Tx constructs were optically transparent (Figures 2A, S2). Cryo-scanning electron microscopy (cryo-SEM) imaging showed that the surfaces and cross sections of both Tx and unmodified implants shared similar surface features of interconnected lamellae (Figure 2B, C). Detailed morphological analyses of the samples was conducted to compare the structural integrity of the Tx and unmodified control implants Fig. 2 (D-F). The inner structural morphology remained conserved in Tx and there were no critical feature differences caused by the incorporation of the ACV-GNPs. Cryo-SEM through a cross-section of a Tx implant shows similar to biosynthetic cornea lamellate morphology.

**Figure 2.**
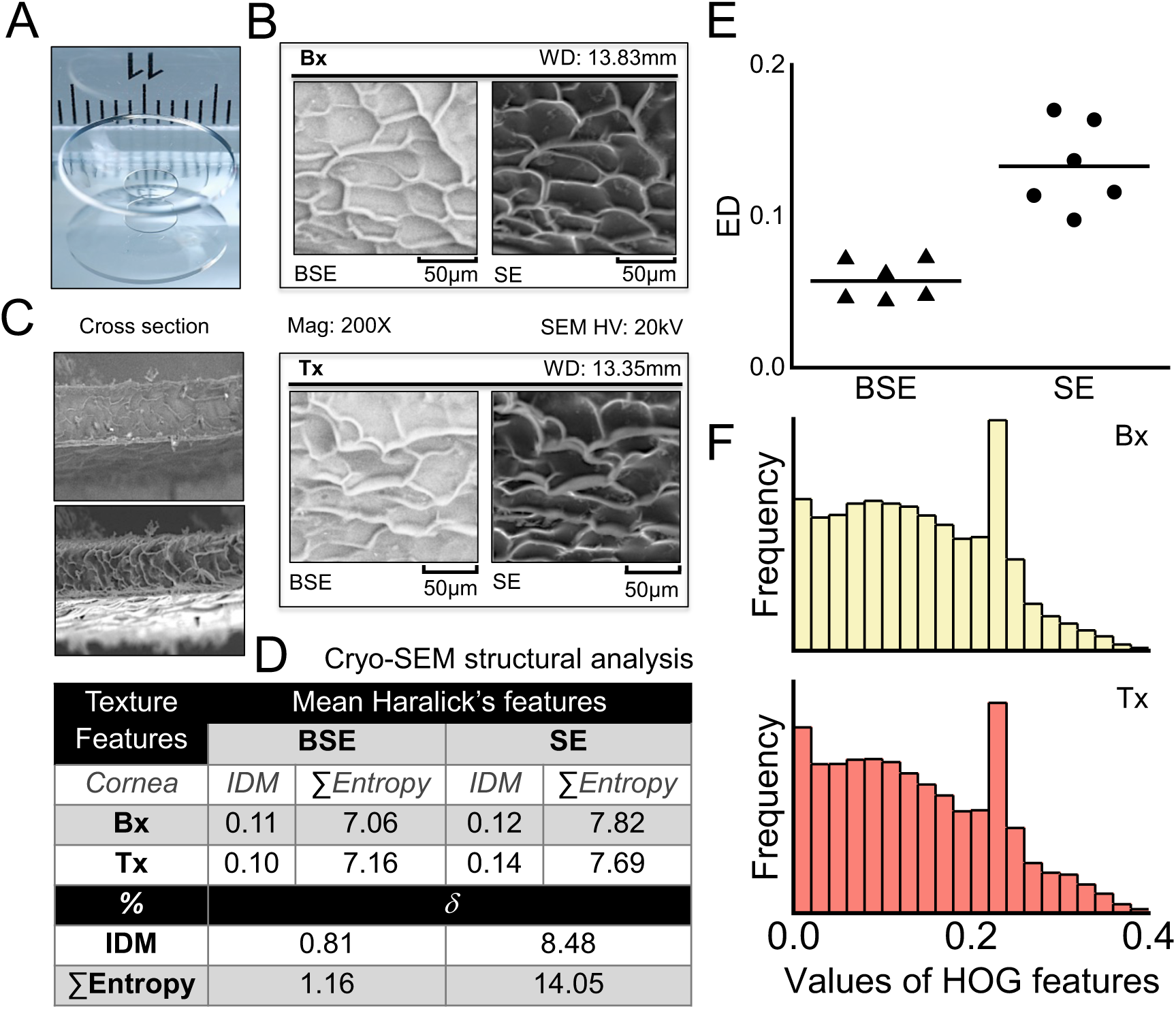
Tx implants. **A**, Physical form of the fabricated corneal implants. **B**, Cryo-Electron Microscopic (EM) features of unmodified control biosynthetic cornea (Bx) and Tx implants and their **C**, 100X cross-sections at 20KV. Backscattered-electron (BSE) images are for comparing compositions, information on topography and crystallinity, whereas secondary electron (SE) imaging are assessment for the topography of the cornea surface (**D-F**) Morphological feature based on HOG and Haralick’s texture to maintain the structural integrity in Tx. Inverse different moment (IDM) is the measure of local homogeneity and sum entropy reflects the measure of non-uniformity in the image during comparison of Bx and Tx electron microscopic features. **E**, Values of Euclidean distance (ED), in each distribution, the central mark is the median. **F**, Values of HOG features.

A detailed histogram of oriented gradients (HOG) analysis together with study and illustration of Haralick’s texture feature analysis show similar local homogeneity and non-uniformity in both Tx and control unmodified hydrogels^25,26^(*detailed in supplementary section 2*). The summary table shows mostly identical physical features. Therefore, the integration of ACV-GNPs into collagen implants did not result in any significant structural perturbations in the overall nanoscale structure of the collagen lamellae.

### Physical and biocompatibility properties of Tx implants

Rheological examination of implants showed that storage the modulus of the Tx was higher than those of controls (Bx), indicating higher stiffness after incorporation of ACV-GNPs (Fig. S8a). The loss modulus (G″) represents the energy dissipated during shear, which gives an indication of the ability of a material to disperse mechanical energy through internal molecular motions, i.e. the viscous response of a material. The viscosity of the Tx was higher than that of controls. When G′ > G″, the hydrogel behaves more like an elastic solid, which was true for both implants (*Supplementary section 4*). Transmittance of white light was comparable between the two sets of formulations (Fig. S8b). Although, the % transmittance was slightly higher in the Tx cornea compared to controls, the increment is almost negligible. This observation again indicates that integration of the nanosystem into collagen implants did not result in transmission changes. Backscatter values for both Tx and control implants were also similar (Fig. S8c).

The thermal stability of Tx implants examined and confirmed by differential scanning calorimetry (DSC). The thermal transition for the control collagen implant was statistically significant (p ≤ 0.05) with the Tx implants (Fig. S8C). GFP-expressing human corneal epithelial cells (GFP-HCECs) seeded onto 3D composite Tx corneas proliferated at a higher rate over the five-day observation period than on control implants (Fig. S8d-e). The incorporation of GNPs or ACV-GNPs had no observable cytotoxic effects and instead resulted in promoted proliferation (by more than 50% at 48h) when compared to unmodified implants (Figure S8d-e). Using different hydrodynamic sizes of GNPs, we also showed that GNPs can promote proliferation of HCECs (*supplementary section 5*) to different extents and this offers flexibility in selecting the GNPs^27,28^ for custom-tuned Tx implants.

### Antiviral activity of Tx implants

The antiviral activity of the Tx implants compared to unmodified controls and Tx implants containing ACV -GNPs was assessed using a viral focus forming assay (FFA) that allows for direct determination of antiviral activity. Viral proteins expressed by infected cells were detected using fluorescently-labelled antibodies, providing a more sensitive assay than the traditional plaque assay. Labelled infection foci are shown in Fig. 3 Here, a HSV 1/2 polyclonal antibody conjugated to FITC was used to stain HSV-1 transient exposure-infected confluent cells growing on the implant surface. Fig. 3A shows the % focus forming units (FFU) present within cells cultured on collagen hydrogels (only negative controls, implants loaded with ACV only, and Tx with ACV-GNPs).

**Figure 3.**
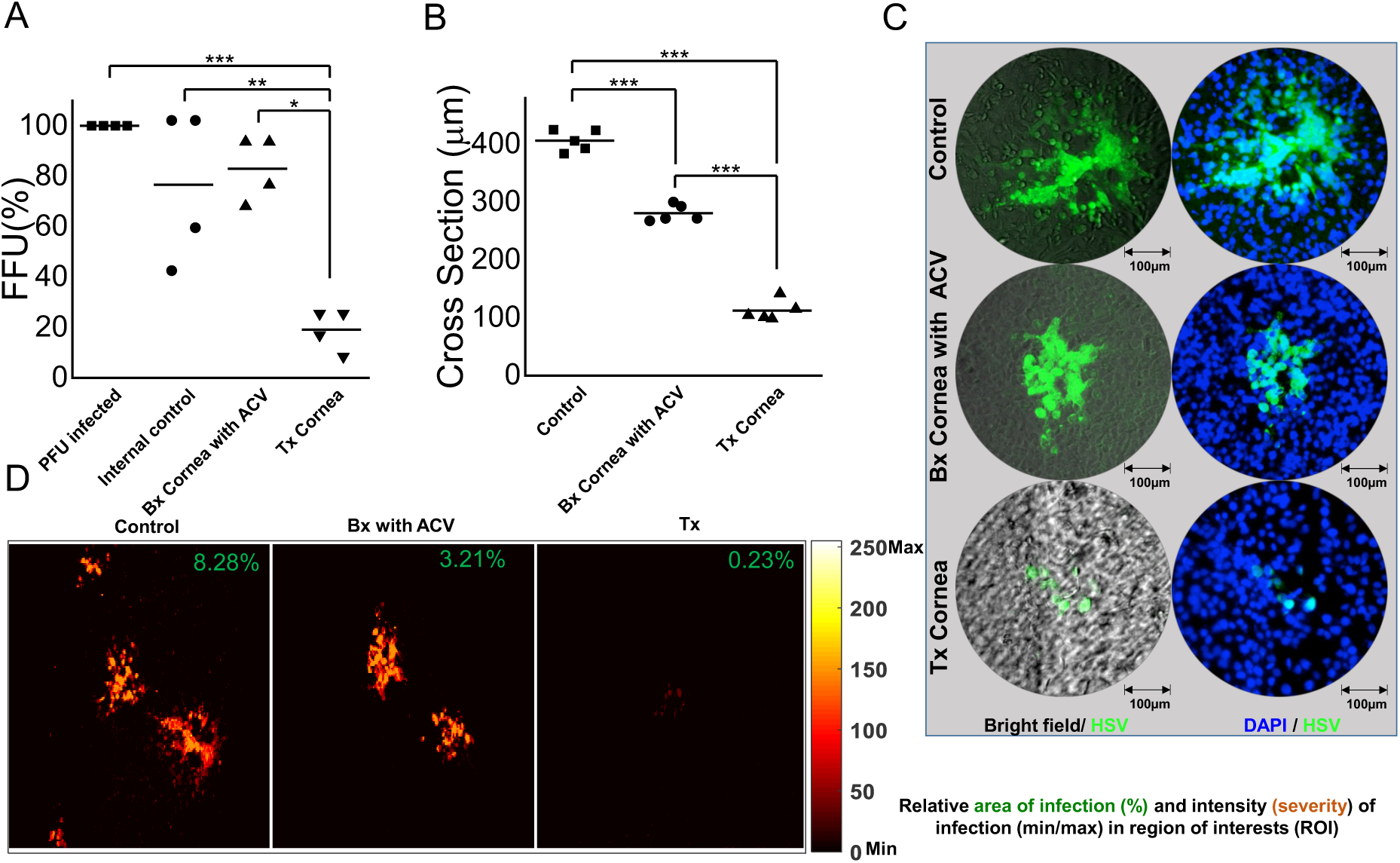
*In vitro* Anti-HSV activity of Tx implants. **A**, Comparative HSV infection measured using Focus forming assay (FFA) and expressed as % FFU (Focus Forming Unit) for direct determination of antiviral activity on the infected Tx. **B**, Relative infection cross section diameter (µm). **C**, Immune-stained individual virus infected focus using anti-HSV specific antibodies**. D**, Severity of the infection represented at Region of Interest (ROI) scaled image after HSV focus formation

A marked reduction in the number of infected foci on Tx was observed compared to control implants with ACV only (p ≤ 0.001). In addition, a significant reduction in the size of infected cell foci on Tx compared to controls was observed (p ≤ 0.0001). The FFA was specifically useful for quantifying the infection on a layer of human epithelial cells growing on three-dimensional, 500-µm thick corneal implants (Fig. 4C). Furthermore, a comparative quantification performed in a region of interest (ROI) (Fig. 4D) by mapping the minimum and maximum intensity of the infection showed that on Tx, not only the number and size of infection foci, but also the relative intensity of fluorescence of the infected foci were greatly reduced compared with the controls.

**Figure 4.**
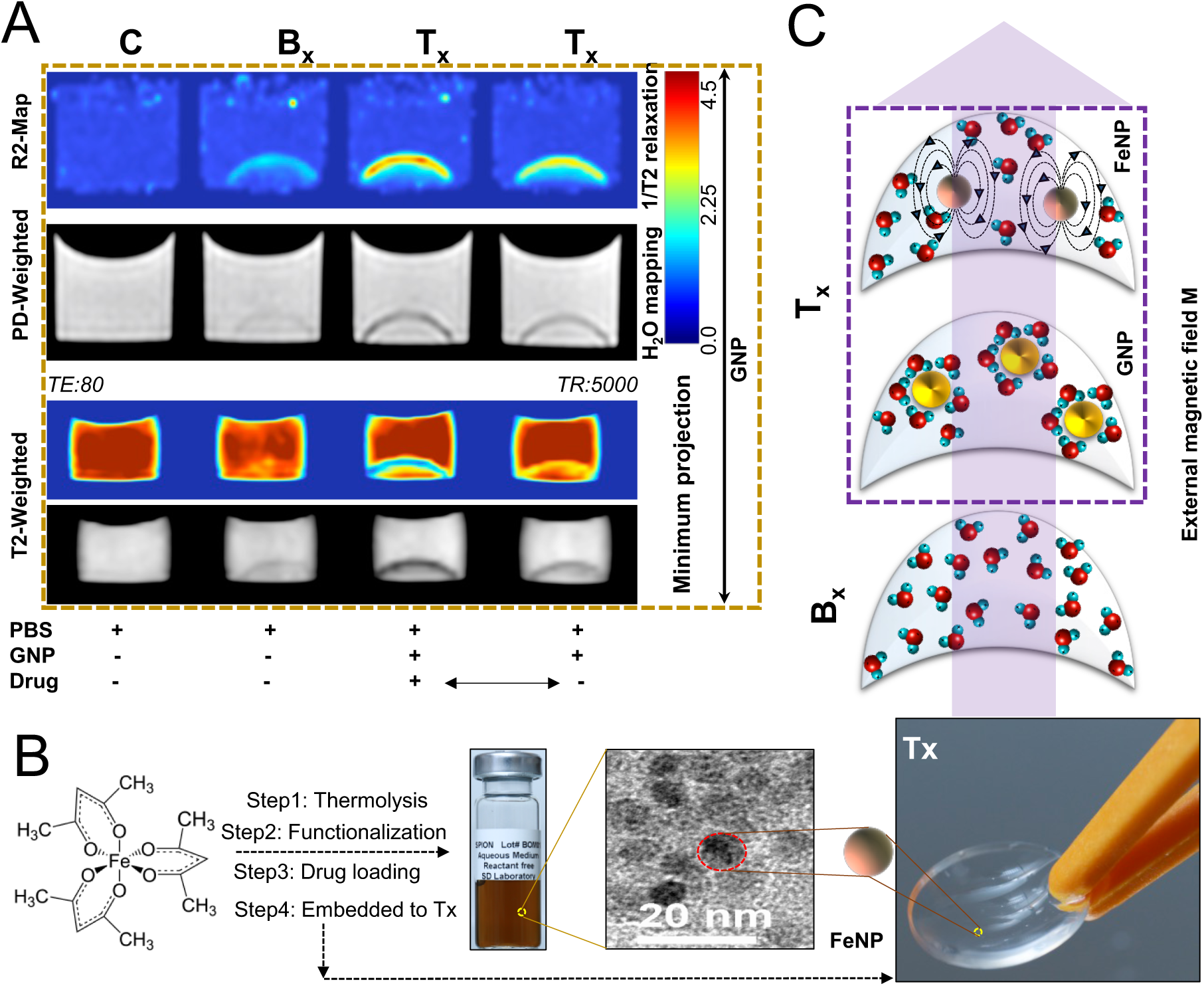
Theranostic corneal implant with non-invasive. **A**, MRI contrast using GNPs based nanosystems for transverse relaxation rate (R2 Map) and proton density weighted (PD-weighted) water mapping. Minimum projection of the transversal relaxation time T2-weighted MRI images in presence and absence of the loaded drugs (bottom panel). **B**, Schematic representation of the MRI-enabled iron based nanosystem developed for Tx **C**, Schematic illustration of the contrast effect influenced by the magnetic relaxation process of the protons in water molecules^29^ of the nanosystems associated with Tx in the presence of external magnetic field during MRI.

### Real time MRI monitoring of drug release from Tx implants

Non-invasive MRI assessments of the Tx microenvironment and tracking the drug molecules in the transplanted implant would be a dramatic step toward corneal tissue regeneration procedure. Such microenvironment and the associated role player interactions with the other eye compartments determine the regenerative fate of transplanted cornea and are important in monitoring individual disease prognosis. However, there are few techniques that can assess these properties non-invasively, globally, in real time, and quantitatively. Implants containing FeNPs were developed to allow the use of multi-modal MRI to track the corneal water environment and the amount of drugs present. While GNPs based Tx implants provided sufficient contrast by both transverse relaxation (T2) and the water content sensitive proton density (PD)^29^ (Fig. 4A) are not sufficient enough to precisely monitor associated water environment. Fig. 4B is a schematic that illustrates the possible interactions of nanosystems for ordering of water molecules that may account for mapping observation. However, the mapping of the extent of drug release is only possible with the enhanced contrast ability of the implants.

To overcome issues of mapping within the human cornea, an FeNP-based nanosystem for ACV delivery was developed. Fig. 4B, and the associated *Supplementary section 6*, shows the steps in the incorporation of ACV-loaded FeNPs (ACV-FeNP) into these Tx implants.

The resulting FeNP based Tx implants when tracked using MRI, showed changing profiles that are related to the extent of drug present (Fig. 5A). Detailed examination of the changing profiles by T2 and PD weighted images showed that the changes in MRI contrast could be quantitatively correlated to the associated water environment and the amount of intra-corneal drug molecules present in a concentration dependent manner (as demonstrated in Fig. 5A). This FeNP based Tx allow us to surveil the drug release profile by changing the MRI contrast R2 and successfully monitored for 60 days (Fig. 5B) including different initial drug dose (5C).

**Figure 5.**
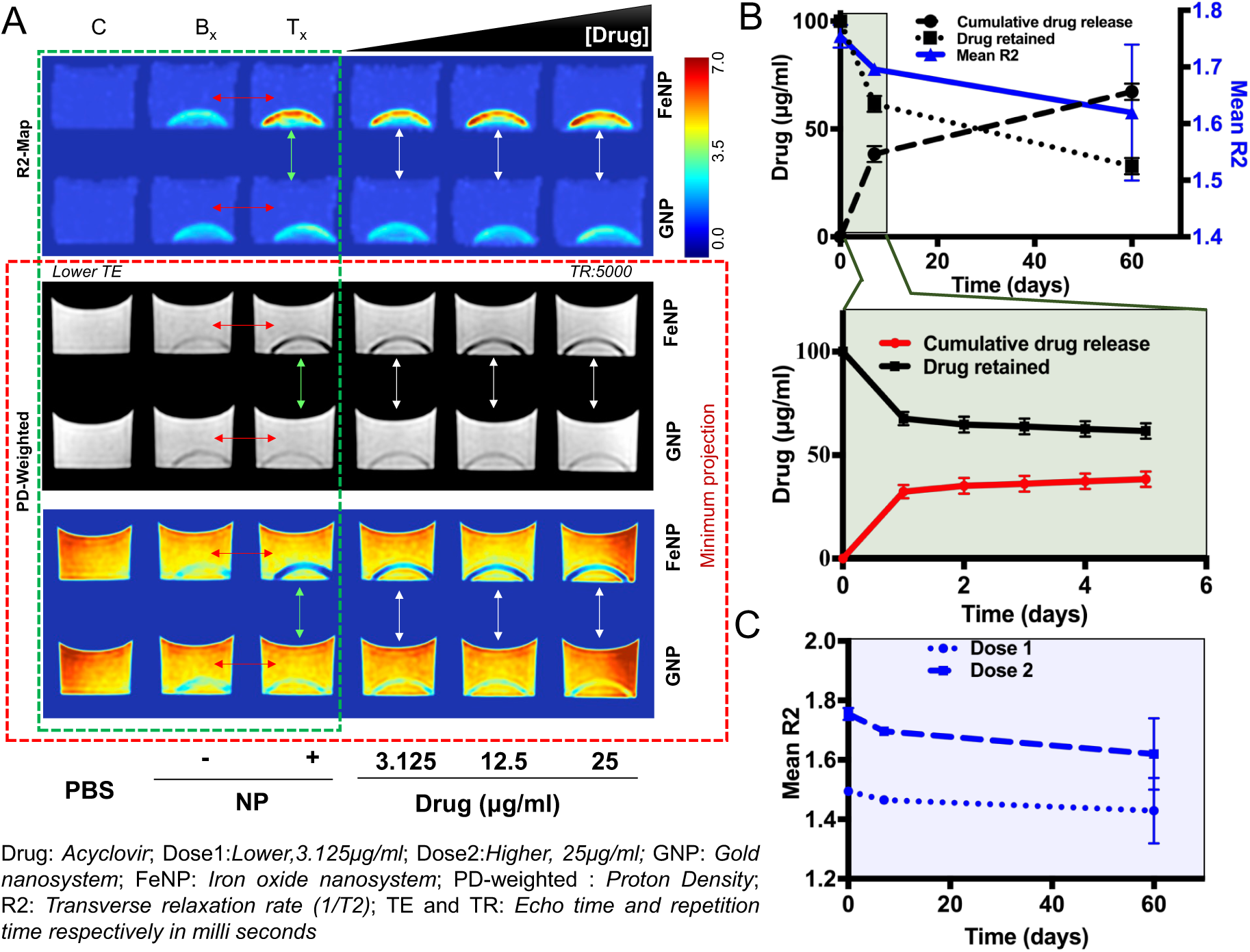
Tx implant for non-invasive monitoring. **A**, Quantitative and comparative MRI mapping of superparamagnetic iron oxide based nanosystem for development of enhanced MRI contrast Tx. Comparative R2 contrast in presence and absence of nanosystem indicated by red arrow (green box) and comparative MRI contrast on the extent of drugs present within the Tx indicated with white arrow. Local water environment can be observed with minimum projection in both GNPs and FeNPs based Tx implants for proton density weighted (PD-weighted) images and can be helpful indicator for infection, inflammation, integration and regeneration process of the Tx. **B**, Extended cumulative release and retention profile of the antiviral drug (ACV) for two months **C**, Relative magnetic resonance imaging MRI contrast R2 (follow-up of the effective transverse relaxation rate) with two different dose of loaded drug in Tx.

## Discussion

A simple theranostics toolbox has been developed for pro-regenerative corneal implants with functionalities suitable for an effective alternative to human donor cornea. A theranostic approach is needed for patients at high risk of rejecting implants in a model system such as with HSK, where viral reactivation is very well documented. The ideal theranostic approach should consider patient safety and selectivity for the site of action, and contain the ability to diagnose disease conditions and to provide efficient delivery of therapeutics agents. The diagnostic component required to track viral reactivation was developed earlier by detection of a cytokine with a major role in activation of the inflammatory cascade associated with disease recurrence^17^. Current results show that it is possible to integrate the therapeutic component of the implantation-cum-monitoring theranostic package into the implants. As such we have selected components that will allow for safe clearance from the body as it is or as a degraded nontoxic product, or integration within the body^30,31^.

As a backbone of this work, we used the formulation of the biosynthetic cornea that has been shown to promote effective regeneration of corneal tissue and nerves in a mini-pig model^24^. ACV, a widely used antiviral agent was selected as this or a variant is likely to be the drug of choice in treatment of patients with HSK. Being hydrophobic and highly insoluble in aqueous solutions, ACV offers very limited opportunity to modify and retain its functional activity^32^ to inhibit viral DNA polymerase enzyme to stop viral replication. ACV also has poor solubility in aqueous solutions and low permeability^33^ that makes difficult to formulate prophylaxis^34^. Direct incorporation of ACV within collagen hydrogels resulted in burst release and rapid diffusion of drug out of the polymer matrix^35^ GNPs were therefore used as a carriers for ACV, and the ACV-GNPs were then incorporated into the collagen-based implants. Our results showed that GNPs can be loaded with ACV within the zwitterionic range of the drug. The hydrodynamic diameter of the drug loaded particles increased from 20 to 81 nm. This was the case in our Tx where there were no changes to the optical transparency after loading of ACV-GNPs. We have previously released ACV from silica nanoparticles^36^ and release from liposomes have also been described previously. In this study, we tethered ACV to GNPs for delivery. Once released, the ACV was fully potent and able to stop HSV-1 activity. Furthermore, the ACV had no adverse effects on cell proliferation at the doses used and in fact, proliferation rates were enhanced in the presence of GNPs. The proliferation rate of HCECs was higher, which could be a feature of a mechanically stiffer hydrogels or confirmation of reports that the presence of GNPs can promote cell growth^27^.

The regeneration of corneal tissues and nerves can be assessed in real-time by *in vivo* confocal microscopy in animal models^37^ or patients^38^. However, this technique is limited by topographic repeatability in addition to lack of global representation^39^ and is unable to track the amount of drug infused or released. MRI is commonly used in health care centers to obtain detailed images within the body without exposure to radiation with the help of gadolinium and iron-based contrast agents. Here, we showed that FeNPs could be incorporated into Tx implants to probe the extent of drugs release from the implants as monitored by MRI reflected the release of ACV. The capacity to track the local environment through proton density is particularly important during corneal pathogenesis^40,41^. Reactivation of the virus in the implanted eye in the clinical setting can range from few days to months and hence the retention of drugs is important to keep them pre-medicated for prolong period to combat against reactivation. More importantly, this approach can paid way in future therapeutics by releasing recombinant nerve growth factors to address vital issues such as in case of delayed epithelialization. It is also possible that the pattern of contrast in the MRI images will change as neo-tissue is regenerated within the implants, and as the implants are remodeled. The evaluation of Tx and control implants within implanted corneas in animal models, and the diagnostic contact lens previously developed will be an important next step in the integration of theranostic implant and monitoring system for future translation.

## Conclusion

Cell-free, pro-regeneration collagen-based corneal implants incorporating nanosystems releasing ACV have the potential to prevent peri-operational reactivation of HSV-1 viruses in compromised corneas. Together with contact lenses that were previously developed to detect interleukin-1 as an inflammation marker, we demonstrate the feasibility of a two-part theranostic system for use in HSK corneas at high risk of rejection due to disease recurrence. While *in vivo* feasibility testing in animal models is merited, we have demonstrated the feasibility of a theranostic system that could be used to deliver drugs and other bioactive molecules to the cornea and to allow precise monitoring through a combination of MRI and clinically available *in vivo* confocal microscopy. This “nano toolbox” with its precise monitoring capability will have an important role particularly in the pre-clinical safety and efficacy testing of new composite implants with therapeutic functions. Further development of the toolbox will allow fabrication of customised implants for patients.

## Materials and Methods

### Development of the nanosystems

#### Synthesis of GNPs and conjugation with ACV

GNPs were synthesized using Turkevich and Frens method with required modifications^42^. Briefly, an aqueous solution of HAuCl_4_ (Sigma-Aldrich, St Louis, MO, USA; 250µM, 25mL) was heated at 90-95°C, close to boiling condition, while stirred continuously with a magnetic stirrer. Freshly prepared 100mM trisodium citrate dihydrate (Sigma-Aldrich, St Louis, MO, USA) solution was then added at once, in different amounts (final concentrations 1200, 600, 400, 200 and 100µM) depending on the desirable particle size (*Supplementary section 1*). The solution was heated up for additional 25 minutes, resulting in a change of solution colour from pale yellow to orange for smaller particles or purple for bigger ones. After obtaining a persistent color the solution was allowed to cool down to room temperature. The purified stock solution of GNPs was obtained by precipitating and dispersing in ambient medium. For this, 20ml of GNPs solution was centrifuged at 7500 rpm and the supernatant was discarded. This is done to reduce the formation of additional citrate layers between the adsorbed citrate on the surface of the GNPs and the free citrate ions available in the solution and also to remove free Cl- and acetone decarboxylate formed during the course of the reaction process. The centrifuged GNPs were re-suspended in trisodium citrate/citrate buffer and the pH of the re-suspended GNPs colloidal solution was kept above 6.0 for further conjugation process. The purified colloidal suspension of GNPs was kept at 4°C for further use.

Acycloguanosine (synonym of ACV; HPLC grade) was purchased from Sigma-Aldrich (St Louis, MO, USA). Stock solution of ACV was prepared in DMSO (10mg/ml) (Sigma-Aldrich, St Louis, MO, USA). From the purified/concentrated solution of GNPs, 23.78µg/ml of GNPs was used and to this 100µg/ml of ACV was added dropwise. The pH of the purified nanomaterials should be above 6.0, to allow for the conjugation of zwitterionic ampholyte ACV to easily bind to the surface of GNPs through ionic interaction between the negative citrate ions of the surface of the nanoparticles and the basic –NH_2_ groups in ACV.

#### Synthesis of magnetic nanoparticles and conjugation with ACV

FeNPs were synthesized by thermal decomposition of Iron acetylacetonate (Sigma-Aldrich, St Louis, MO, USA). In brief, the dissolved iron acetylacetonate in benzyl ether (Sigma-Aldrich, St Louis, MO, USA) and oleylamine (Sigma-Aldrich, St Louis, MO, USA) mixture was heated at 110 °C degree in nitrogen environment. This mixture was then reheated at 310°C for 2 more hours followed by cooling down to room temperature before it was mixed with 60ml ethanol. The suspension was centrifuged at 3000 rpm for 20 min and the supernatant was discarded and the precipitate was allowed to air-dry before resuspension in 20 ml hexane (Sigma-Aldrich, St Louis, MO, USA). The mixture was then been centrifuged at 25000g for one hour to remove larger particles and the supernatant was collected for functionalization.

Homo-dicarboxylic polyethylene glycol (PEG-2kd) (JenKem, USA), dimethyl formamide (Sigma-Aldrich, St Louis, MO, USA), chloroform (Sigma-Aldrich, St Louis, MO, USA), and N-ethyl-N-(3-dimethyl aminopropyl) carbodiimide hydrochloride (EDC) (Thermo Fisher Scientific, USA) were mixed in a reaction flask. The mixture was then stirred at room temperature for 30 minutes. Next dopamine (Sigma-Aldrich, St Louis, MO, USA) was added to the mixture and stirred at room temperature for 90 minutes followed bythe addition of oleylamine coated synthesized nanoparticles to the reaction vessel and overnight stirring at room temperature. Particles were allowed to settle down and washed with hexane, followed by drying. The particles are suspended in water and were centrifuged 3000 rpm for 20 min to remove any aggregates formed. Then the particle suspension was spun at 2500-3000 rpm for 40 mins in nano-sep (Pall Pvt Ltd, MI, USA) (10 kd) to remove un-reacted PEG, and finally suspended in citrate buffer at around pH 7.5 to allow the binding of the zwitterionic ampholyte ACV to the surface of FeNPs through ionic interaction between the negative carboxyl ions of the surface of the nanoparticles and the basic –NH_2_ groups in ACV.

#### Flocculation test

GNPs and GNPs-ACV were incubated with different concentrations of NaCl solutions (4, 8, 16, 25 and 50 mM) in a 1:1 volume ratio. All the samples were incubated at room temperature for 10 minutes and the hydrodynamic diameter, zeta potential and UV-visible absorbance were measured^43^.

#### Dynamic Light Scattering (DLS), Zeta Potential and UV-Visible absorption measurement

The hydrodynamic diameters (HD) and Zeta potential (ξ) measurements for both bio-conjugated and unconjugated nanomaterials under investigations were performed using a Zeta Sizer Nano ZS90 equipped with a red (633nm) laser (Malvern Instruments Ltd., England). All reported HDs are based on averaged intensity. For each sample, three measurements were conducted with a fixed 10 runs each. The same instrument was used to measure the ξ of the particles in the colloidal suspension by analysing the relative velocity of the particles with respect to the fluid. Again for each sample three measurements were performed with a fixed 10 runs each. All the UV-Vis experiments were conducted using Thermo UV-Visible Spectrophotometer (Lambda 25, PerkinElmer, Akron, OH, USA).

#### Fourier Transformed Infrared Spectroscopy (FTIR)

Surface characteristic changes of GNPs after conjugation of ACV were determined by Attenuated Total Reflectance (ATR-FTIR). The measurements were performed using a PIKE MIRacle ATR accessory with a diamond prism in a vertex 70 Spectrometer (Bruker, Massachusetts, USA) with a DLaTGS detector. The whole system was continuously purged with nitrogen and the IR spectra were acquired at 4cm-1 resolution. A total of 64 scans were performed between 4400-600 cm^−1^.

#### Fabrication of implants

Briefly, 0.5ml (∼500mg) of 18% (wt/wt) porcine collagen solution type 1 (Nippon Meat Packers, Japan) was added in a syringe mixing system as previously described^24^ and buffered with 150 µl of 0.625 M morpholinoethanesulfonic acid buffer (MES, EMD Chemicals, Gibbstown, NJ). Calculated amounts of N-Hydroxysuccinimide (NHS; Sigma-Aldrich, MO, USA) (5% (w/v) in MES) and N-(-3-Dimethylaminopropyl)-N’-ethylcarbodiimide hydrochloride (EDC; Sigma–Aldrich, MO, USA) (5% (w/v) in MES) solutions were added and the reactants were thoroughly mixed at 0°C (EDC: collagen–NH2 (mol:mol) = 0.7:1, EDC:NHS (mol:mol) = 1:0.5). The final mixed solution was immediately cast into cornea-shaped moulds (12 mm diameter, 500 µm thick) or between two glass plates with 500 µm spacers and kept at 100% humidity at room temperature for 24 hrs and then at 37°C for 1 hr. Hydrogels were then carefully remoulded and kept in 10 mM PBS at 4°C. Two different types of collagen corneal implants were prepared (i) Biosynthetic cornea without any nanosystems (Bs) and (ii) nanosystem (GNPs, FeNPs) embedded theranostic cornea (Tx). The final concentrations of the different nanosystems and loading of ACV adjusted accordingly within the collagen matrix are mentioned in Figure 6 (d). In all cases, the additives were added to collagen mixing system before adding EDC and the volume was adjusted to keep the final concentration of collagen at 10% (wt/vol). In the control-without any nanosystems and ACV hydrogel, respective amount of buffer was added to keep the dilution factor the same as that of experimental cornea. This formulation was used in all characterization and *in-vitro* studies unless stated otherwise.

#### Characterization of implants

White light transmission and backscattering of hydrogels were determined at RT on a custom-built instrument^44^. Differential scanning calorimetry (DSC) was performed using a Q2000 differential scanning calorimeter (TA instruments, New Castle, DE) as previously described^45^ to measure the thermal transition and evaluate the degree of cross-linking in the hydrogels. Heating scans were recorded within the range of 8 to 80°C at a scan rate of 5°C min^−1^. Samples weighing in the range of 4 to 7 mg were surface-dried and hermetically sealed in an aluminium pan to prevent water evaporation. A resulting heat flux versus temperature curve was then used to calculate the denaturing temperature (T_d_). Denaturing temperature comes from the T_max_ of the endothermic peak. Mechanical properties of hydrogels were measured using a parallel plate rheometer (AR 2000 rheometer, TA instruments, Inc., UK). Hydrogels were trephined in cylindrical shape (500um thick, 8mm diameter) and storage modulus (G’) and loss modulus (G’’) measurements were recorded at a frequency range of 1-8Hz at 25°C using 8mm aluminium plate geometry. The gap was corrected starting from the sample height and compressing the sample to reach the sample to reach a normal force of 0.3N. The morphology of the biosynthetic hydrogel cornea constructs (with and without nanoparticles and anti-viral drug) was studied using scanning electron microscopy (SEM). Processing of the hydrogel prior to conducting SEM has been performed as described previously^46^. SEM micrographs were obtained at 20kV at various magnifications on a scanning electron microscope (Model S225ON, Hitachi, Japan). Comparisons were drawn against the control Bs (without drug and nanomaterials) and Tx cornea (containing the bio conjugated anti-viral drug and nanoparticles).

#### ACV release study

ACV-GNP containing hydrogels were transferred into a 50ml centrifuge tube containing 15ml 10mM PBS to keep the whole hydrogel immersed in the liquid. The tubes were then placed in a continuous mechanical shaker with 100rpm rotation at 37°C. The buffer in the tubes were collected at day 1, 2, 3, 4 and 5 and substituted with fresh buffer. The absorbance spectra of collected PBS buffers were recorded at 250nm using Thermo UV-Visible Spectrophotometer (Lambda 25, PerkinElmer, Akron, OH, USA) and the cumulative release of ACV from the hydrogel construct was evaluated using a standard ACV curve.

#### In vitro cell adhesion and proliferation assays

To examine the cell attachment and proliferation rate on different surfaces, green fluorescent protein (GFP) transfected, immortalised human corneal epithelial cells (HCECs)^47^ were seeded (5000 cells/well) onto 6mm hydrogel discs in a 96-well plate. The GFP-HCECs were maintained in keratinocyte serum-free medium (KSFM; Life Technologies, Invitrogen, Paisley, UK), supplemented with 50 µg/ml bovine pituitary extract and 5 ng/ml epidermal growth factor, in a 37°C incubator with 5% CO_2_. As controls, cells were also seeded on a regular tissue culture plate (TCP) and on a non-binding surface plate (data not shown). At 4, 24, 48, 72, 96 and 120 hrs, images of three different areas on each hydrogel disc were captured using a fluorescence microscope (AxioVert A1, Carl Zeiss, Gottingen, Germany) for quantitation.

#### Cell viability assay

Cell viability after incubation with the different GNPs and FeNPs concentrations were determined using the MTS assay, which is based on the reduction of MTS (3-(4,5-dimethylthiazol-2-yl)-5- (3-carboxymethoxyphenyl)-2- (4-sulfophenyl)- 2H- tetrazolium) compound by living and metabolically active cells to generate a soluble coloured formazan derivative. HCECs were subcultured in 96-well plates containing 100µl of the culture medium. Twenty-eight thousand HCECs were seeded into each well. The cells were treated for 12, 24 and 48 hrs with GNPs and GNPs-ACV. After that, 20% v/v of MTS reagent in cell culture medium was added and cultures were incubated further for 3 h at 37°C. Absorbance was measured at 490 nm and 650 nm as a reference using a spectrophotometric microplate reader (VERSA Max Microplate Reader, Molecular Device, Sunnyvale, CA, USA). Each sample was tested in triplicates and the average survival percentage was calculated with respect to the control (no GNPs/GNPs-ACV added). Before the addition of the MTS reagent, microscopic imaging (AxioVert A1, Carl Zeiss, Gottingen, Germany) was performed to assess the viability of the cells.

#### In-vitro fluorescent focal assay (FFA) for viral count

The antiviral property of the hydrogel containing GNPs with/without ACV was evaluated. Hydrogels were cut at 6mm diameter and placed in a 96-well plates. Forty thousand HCECs were cultured on these hydrogels until they were confluent. Cells were then infected for one hour at 37°C with the HSV-1, strain F (a gift from Earl Brown, University of Ottawa, Ontario, Canada) at a concentration of 6 plaque forming units (PFU) per each well to avoid overlap and distinctly observe individual focus. A negative control was used containing uninfected cells and a positive control containing cells infected with the virus and without GNPs. Each different treatment was tested in triplicate. The cells were washed in phosphate-buffered saline (PBS; ThermoFisher Scientific, Waltham, Massachusetts, USA) and 100µl of new medium were added to each well. The infected monolayer was incubated at 37°C in 5% CO_2_ until the observation of plaques (approximately 24 h). The cells were then washed in cold PBS and fixed overnight in ice-cold absolute methanol. Next day the cells were washed twice with PBS in RT and incubated with HSV 1/2 polyclonal antibody FITC conjugate (ThermoFisher Scientific) at a dilution of 1:100 in PBS for 3hrs at RT and then washed twice in PBS. The cells were mounted in 1:1 glycerol-PBS and imaged using a fluorescence microscope (Leica DMi8, Leica, Wetzlar, Germany). Fluorescent focus units (FFU) located in each well were counted and virus titers were calculated and expressed as percentage (%) FFU with respect to control.

#### Statistical analysis

Unpaired t-test was performed to compare each different intervention with the control. Values are represented as mean ± S.D. A p value<0.05 wasconsidered statistically significant. All the tests were performed on MATLAB analysis software (MATLAB 2016B, Mathworks, USA).

#### Magnetic Resonance Imaging and Analysis

All MR-images were acquired with a 32 channel head coil in Philips Ingenia 3.0T scanner (Royal Philips, Amsterdam, Netherlands). Proton density weighted images were acquired using a multi-slice turbo spin-echo sequence, TR 6 sec, effective TE 8 ms, flip angle 90°, bandwidth 348 Hz, acquired in-plane resolution 0.6×0.6 mm^2^, slice thickness 0.8 mm and a turbo factor of 15. T2-weighted images and R2-maps (1/T2) were generated on the scanner based on a multi-echo turbo spin-echo sequence with 20 echoes, first TE 10 ms, delta-TE 10 ms, TR 5 sec, flip angle 90°, bandwidth 177 Hz, acquired in-plane resolution 0.6×0.6 mm^2^, slice thickness 1.3 mm and a turbo factor of 20. Minimum intensity images were generated using OsiriX (Pixmeo SARL, Geneva, Switzerland).

## Supporting information

Supplementary section

## Author’s contribution

HP conceived, designed, performed experiments and supervised the project, MA, GZ and GP. HP, GP, MA and MI have developed the nano toolbox and Tx. SD and HP have developed the cornea compatible MRI nanosystem for theranostics. HP, TR and MA designed, optimized and analysed the magnetic measurements. HP and AD have analysed the electron microscopy experiment. AG and MJ helped in Bx and Tx spectroscopy (uv-vis, fluorescence and IR) and analysis of the data. JH supervised MA and gives feedback during development of the Tx. MR, JH, NS helped with clinical significance, scientific discussions, critical comments and intellectual inputs. HP, MA, MI, and GP drafted the MS. We confirm that the manuscript has been read and approved by all named authors and that there are no other persons who satisfied the criteria for authorship but are not listed.

## Acknowledgement

We cordially thank Emilio Alarcon from University of Ottawa, Canada for helping with electron miscopy experiment and May Griffith from Linkoping University, Sweden for providing research materials and laboratory set up. This work was supported by EU H2020 Marie Sklodowska-Curie Individual Fellowship (Grant no: 706694), MIIC Strategic Postdoc Grant and MIIC Seed Grant at Linkoping University (LiU), Sweden. We thank Magnus Borga for providing us the magnetic measurement facilities at CMIV, LiU and for his helpful inputs before designing the magnetic study.

