## Supplementary section for "Rational nanotoolbox with theranostic potential for medicated pro-regenerative corneal implants"

### Supplementary Information

***Section1: Nanoparticle toolbox for ACV conjugation and development of Tx implants***

GNPs were synthesized by wet-chemical reduction method using tri-sodium citrate as a reducing and capping agent (S1a). The UV-Vis absorption spectra of the different GNPs (GNP1-GNP5) demonstrated a peak absorbance that varied between 517-551nm (S1b).

**a** $6Ctr^{3-} + 26AuCl_4^{-} + 24H_2O \xrightarrow{90^{\circ}C} 26Au + 6CH_2O + 3HCOOH + 27CO_2 + 104Cl^{-} + 60H^{+}$

**b**

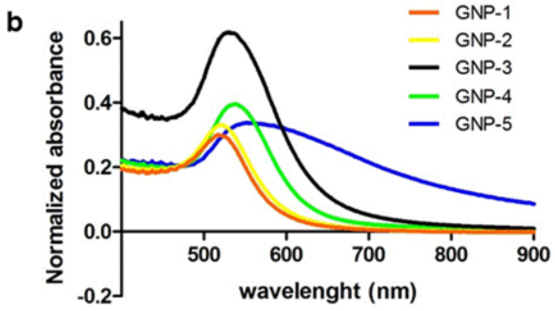


| NP name | SPR max (nm) | Citrate concentration (μM) |
|---|---|---|
| GNP-1 | 517 | 1200 |
| GNP-2 | 520 | 600 |
| GNP-3 | 530 | 400 |
| GNP-4 | 539 | 200 |
| GNP-5 | 551 | 100 |

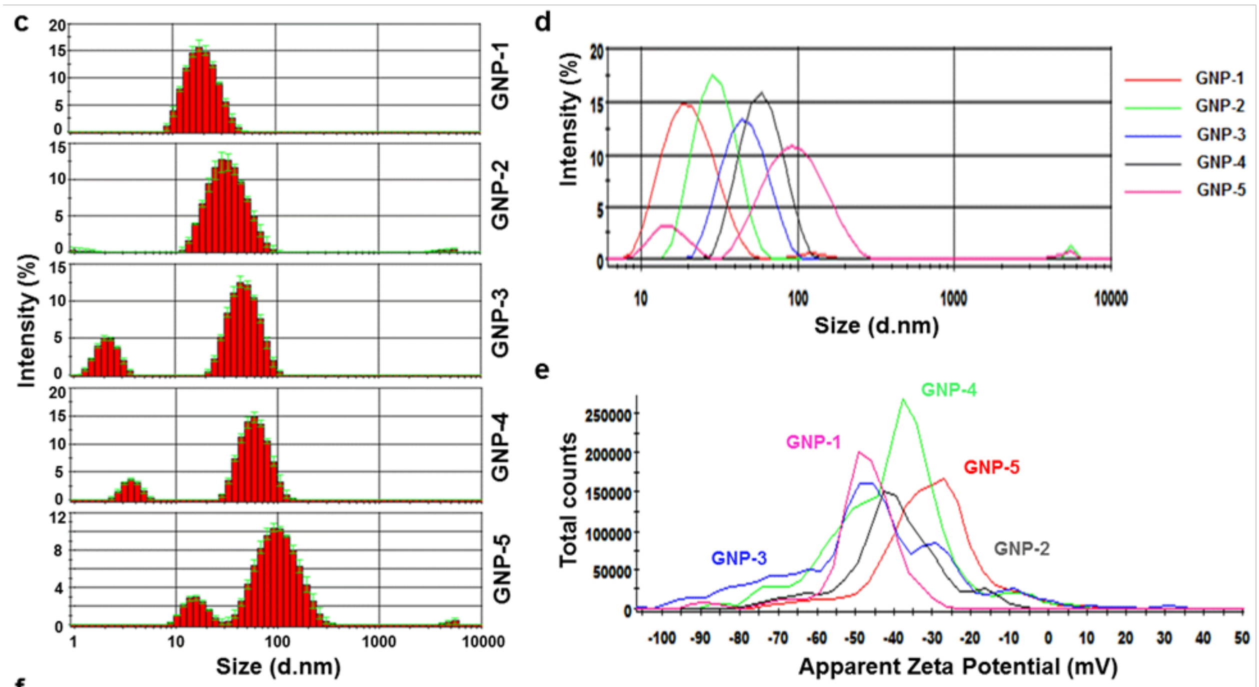


**f**

| NP name | Average size (d. nm) | Mean peak 1 intensity (d. nm) | PDI | zeta potential (mV) |
|---|---|---|---|---|
| GNP-1 | 18.17 | 21.23 | 0.152 | -49.3 |
| GNP-2 | 32.67 | 30.74 | 0.232 | -42.8 |
| GNP-3 | 43.82 | 47.53 | 0.967 | -45.7 |
| GNP-4 | 58.77 | 60.6 | 0.697 | -37.5 |
| GNP-5 | 105.7 | 102.7 | 0.459 | -27 |

**Figure S1**: Summary of the synthesis of different GNPs: **a** Citrate reduction of aqueous auric chloride solution **b** Surface Plasmon absorption of different GNPs **c** Hydrodynamic diameter and the statistical size distribution of the synthesized GNPs **d** Comparative Dynamic Light Scattering (DLS) profile of the synthesized GNPs **e** Relative zeta potential and **f** summary table of all the GNPs

The shift of the Plasmon Resonance absorption peak to higher wavelength with decreasing citrate concentration indicates an increase in size and changes on Surface Plasmon's. The study of DLS indicated the hydrodynamic size distribution of the different GNPs with their frequency distribution ranging from 18nm for the smaller particles (GNP1) to 110nm for the largest ones (GNP5) (Figure S1c). The comparative distribution indicated the mean intensity of the major peak as shown in (*Supplementary Figure S1d*). The zeta potential showed a gradual decrease in the surface negative charge, from -52.5mV to -29.6mV, with decreasing citrate concentration (Figure S1e). A higher zeta potential indicates stronger electrostatic repulsion between adjacent particles and a more stable system, which is very important in the later stage of conjugation and integration into the hydrogel. Table f (Figure S1) summarizes the hydrodynamic properties of the synthesized GNPs (GNP1-GNP5) along with the absorption maxima and concentration of citrate used.

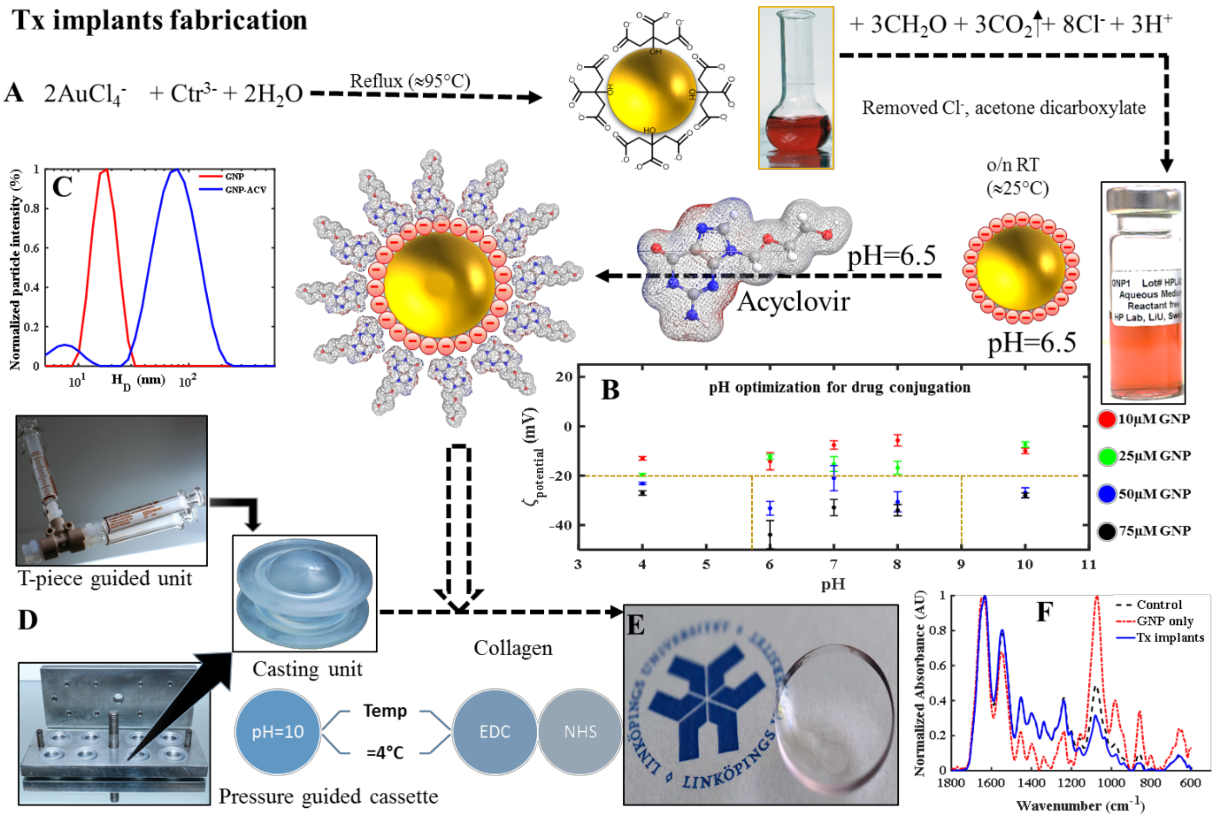


**Figure S2.** Schematic workflow for **A-C** development of Tx using gold nanosystem with optimized conditions. In the lower panel **D** Tx fabrication strategy using T-piece guided mixing and pressure guided jigs with 500µm molds casting unit described with their respective **E** physical form and **F** molecular signatures. Detailed chemical signature and conjugations are described in *supplementary section 3*.

We have found that the synthesized GNPs are biocompatible to human corneal epithelial cells (HCEC) (*supplementary section 5 and Figure S8)* and therefore theoretically any of them can be used. However, we think that GNP-1 to GNP-3 are suitable for ACV conjugation. The colloidal stability of GNP-4 and GNP-5 could limit the scope due to their higher hydrodynamic diameter and effective size-zeta criticality[1]. We have used GNP-1 as our candidate for the present report. The fabrication set-up and workflow for Tx implant formulation is illustrated in Figure S2.

#### Section 2: Quantification of morphological changes between control and Tx implants

*Methodology*

Scanning electron microscopy (SEM) is extensively used to study structural details on the surface of biological samples. The Control and Samples images are compared based on texture analysis. We have cropped several patches of size 200×200 pixels from sample and control images for comparison. Patches created from BSE and SE is shown in Figure S3 and Figure S4, respectively. The comparative study was performed using the patches of samples and control images from BSE and SE, separately.

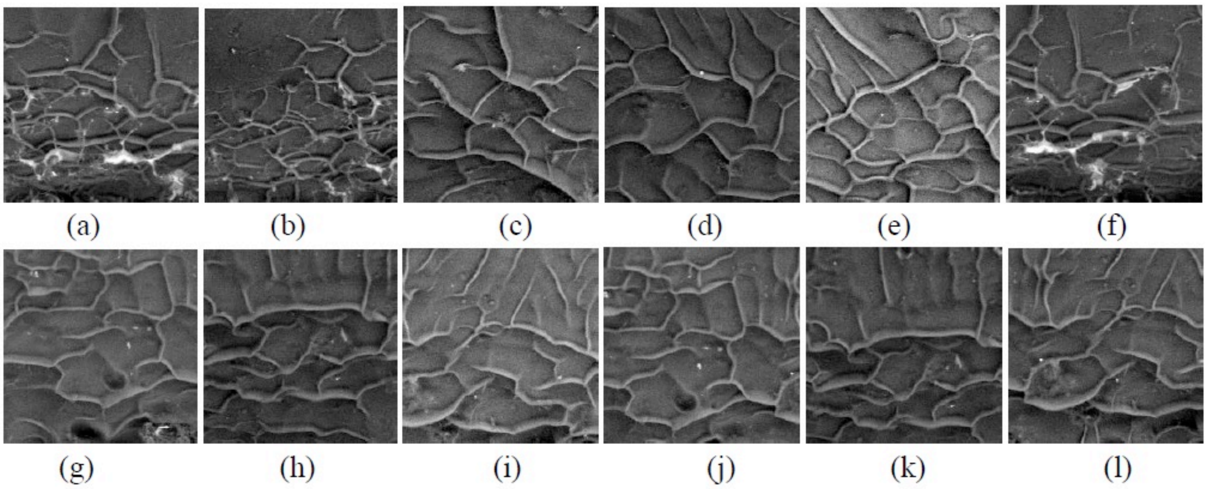


**Figure S3**. **a-f** patches of control and **g-l** patches of samples in BSE

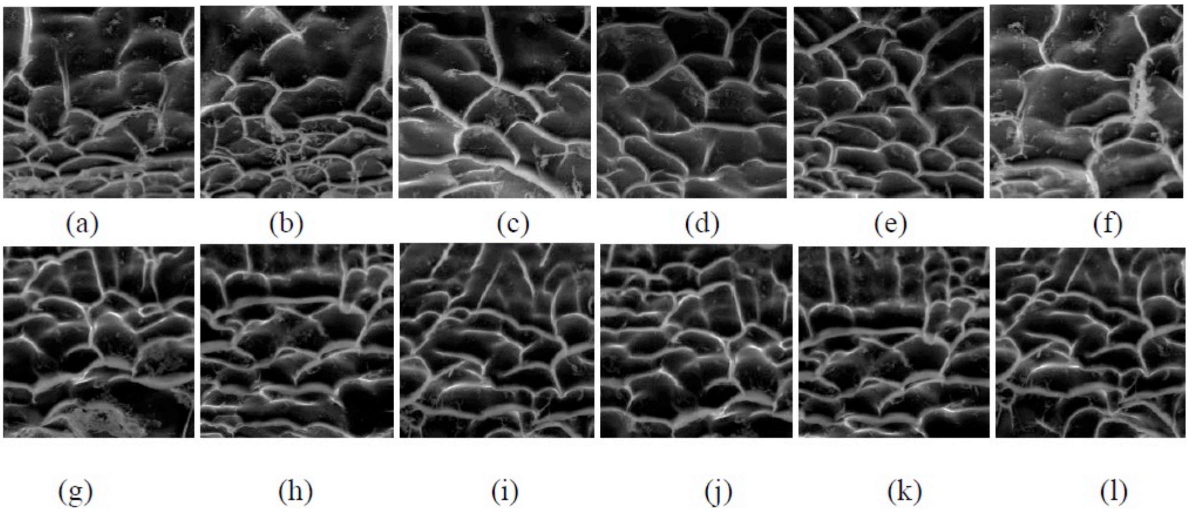


**Figure S4. a-f** patches of control and **g-l** patches of samples in SE

*Morphological change analysis based on Haralick's texture features*

Haralick's texture features[2] are the most commonly used measures of texture, which are computed from the gray-level co-occurrence matrix (GLCM). GLCM is formulated by counting the occurrences of different combinations of gray levels of pixels, separated by a specified distance l and a specified angle $\psi$. The $(i,j)^{th}$ element of a normalized GLCM $p_{(l,\psi)}(i,j)$ with distance l and a specified angle $\psi$ is defined as the probability of occurrence of the pair of gray levels (i, j) at a distance l and at an angle $\psi$. Mathematically, it can be represented as

$$p_{(l,\psi)}(i,j) = \frac{1}{N_x N_y} \mid \{(x_1,y_1),(x_2,y_2) \in (N_x \times N_y) \mid I_q(x_1,y_1) = i,; I_q(x_2,y_2) = j;$$

$$\max(\mid x_2 - x_1 \mid, \mid y_2 - y_1 \mid = l; \angle[(x_1,y_1),(x_2,y_2)] = \psi\}$$

where, $N_x \times N_y$ is the size of the image under consideration, I(x, y) represents the intensity at a pixel (x, y), and Iq(x, y) is the corresponding quantized intensity value with quantization level Nq. Inverse different moment, a measure of local homogeneity and sum entropy, a measure of non-uniformity in the image are used for comparison of sample and control images. Considering p(I, j) = $(i,j)^{th}$ element of the GLCM matrix, inverse different moment and sum entropy can be computed from the set of sample and control patches and taken average over number of patches. Percentage change of mean inverse different moment and mean sum entropy is used to indicate the quantitative analysis as shown in Table-1. Percentage change is defined as

Percentage change = Value of a feature in (Control image - sample image) / Value of the feature in the control image. Percentage change of inverse different moment and sum entropy based on 12 set of patches are provided in Table-II.

Table-1 Definition of inverse different moment and sum entropy

| Feature no. | Feature name | Description |
|---|---|---|
| 1 | Inverse different moment | $\sum_{i=0}^{N_q-1} \sum_{j=0}^{N_q-1} \frac{1}{1+(i-j)^2} p\ (i,j)$ |
| 2 | Sum entropy | $H^2 = -\sum_{k=0}^{2(N_q-1)} p_{x+y}(k) log_2[p_{x+y}(k)]$ |

Table II: Percentage change of inverse different moment and sum entropy based on 12 set of patches

| Image type | Inverse difference moment (%) | Sum entropy (%) |
|---|---|---|
| BSE | 0.81 | 1.16 |
| SE | 8.48 | 14.05 |

*Morphological change analysis based on Histogram of oriented gradient (HOG)*

HOG features[3] capture edge or gradient structure that is an important characteristic of local shape. It does not affect by local geometric and photometric transformations: translations or rotations make little difference if they are much smaller that the local spatial or orientation bin size. The basic idea is that local object appearance and shape can often be characterized rather well by the distribution of local intensity gradients or edge directions, even without precise knowledge of the corresponding gradient or edge positions. In practice this is implemented by dividing the image window into small spatial regions. Over the pixels of the cell, a local 1-D histogram of gradient directions or edge orientations is computed. The combined histogram is obtained by concatenating all local histogram.

Orientation of gradient are visualized for an example sample and control images in Figure S4. Values of HOG features for sample of Figure S5 (a) and control of Figure S5 (b) are shown in Fig. S6 (a) and Fig. S6 (b). The plot depicts that normalized histogram of gradients features are having similar profile for both example sample and control images. Euclidean distance is computed for the feature sets obtained from control and sample set of cropped BSE and SE images, respectively. Box plot of Euclidean distance is shown in Fig. S6 and results indicate small difference in feature space.

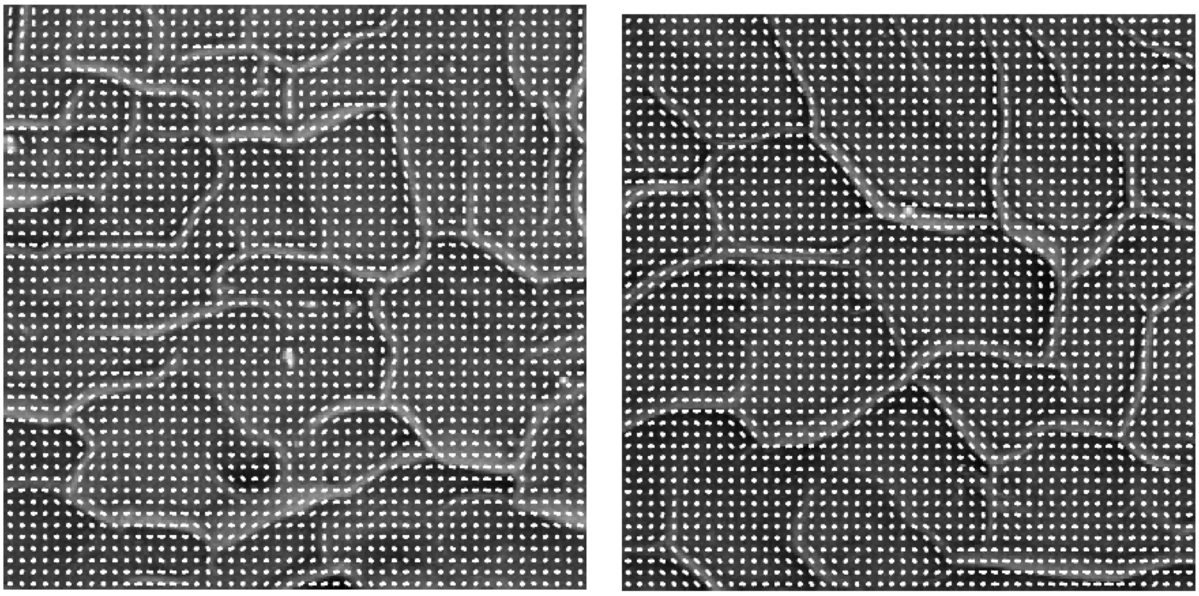

A B

**Figure S5**. **A** Orientation of gradient in Tx implant with respect to **B** control sample for BSE

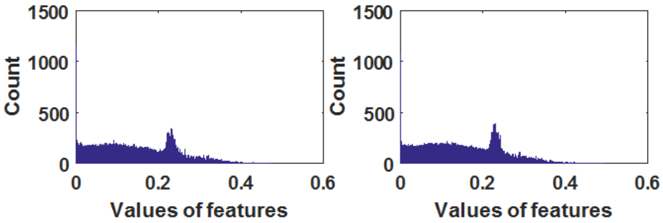


**Figure S6**. Histogram of values of HOG features for sample of Figure S5 a (left) and control Figure S5b (right)

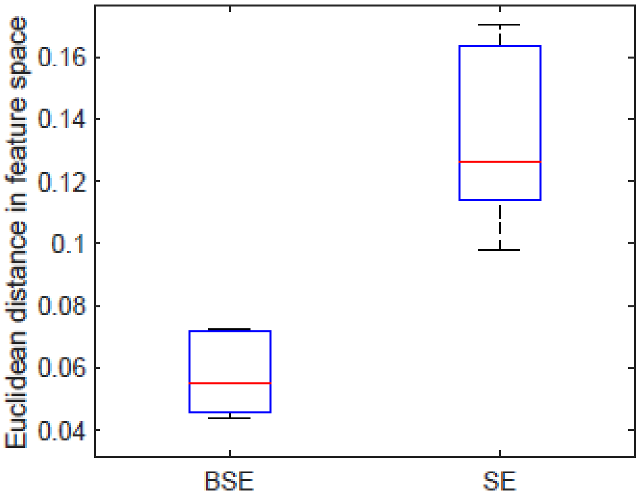


**Figure S7**. Elaborated Box plot of Euclidean distance in feature space as described in figure 2h. In each box, the central mark is the median, the edges of the box are the 25th and 75th percentiles.

*Summary remarks*

Following are the remarks: (i) Haralick's texture feature[2] analysis reveal that local homogeneity and non-uniformity in the Tx implants and control image patches are similar and (ii) Small difference of HOG features of sample image set and control image set, indicating similarity in structural features in image content.

#### Section 3: Fourier Transformed Infrared Spectroscopy

*GNP – ACV conjugation study:* The primary principle of FT-IR spectroscopy is to detect or probe differences on the molecular level arising from changes between the intra and intermolecular interactions[4]. In this study, we have performed ATR-mode FT-IR for demonstrating the intermolecular changes as a result of conjugation between GNP with ACV. In the spectral range from 1300-1000 $cm^{-1}$, where the C-O vibrations are located, there is marked difference in the number of peaks[5]. In case of ACV, there are 10 peaks whereas the bio conjugated GNP-ACV has only a single in this specified spectral region. In the spectral region 1700-1450 $cm^{-1}$, the GNP1-ACV conjugate has only one peak at 1638 $cm^{-1}$ as opposed to the unconjugated ACV which has 6 different peaks. This difference can be ascribed to alterations of C-C, carbonyl, and C-N stretching frequencies[6]. Remarkably, in the region 3600-3200 $cm^{-1}$, the bio conjugated GNP-ACV depicts additional FT-IR spectral absorption peak which are very distinct as shown in table III.

**Table III: Summary of peak positions of the FT-IR spectra of ACV and GNP-ACV**

| Spectral Range ($cm^{-1}$) | ACV | GNP1-ACV |
|---|---|---|
| 1300-1000 | Ten peaks at 1011, 1032, 1047, 1080, 1103, 1180, 1215, 1242, 1281 and 1308 | 1011 |
| 1700-1450 | Six peaks at 1483, 1541, 1576, 1611, 1630 and 1703 | 1638 |
| 3600-3200 | Four peaks at 3291, 3437, 3468 and 3516 | Fifteen peaks at 3208, 3223, 3237, 3246, 3296, 3302, 3318, 3337, 3364, 3379, 3402, 3420, 3466, 3512 and 3584 |

*Tx study:* Structural analysis of the control and Tx constructs were done using Attenuated Total Reflectance FT-IR (ATR-FT-IR) on dry solid samples for both nanomaterials and collagen hydrogel constructs. All the experiments were performed using Bruker Vertex 70 spectrometer equipped with a wide range of accessories for ATR. Scans were conducted from the spectra range of 4000-600 $cm^{-1}$ with 72 repetitions averaged for individual spectrum. The standard spectral resolution of better than 0.4 $cm^{-1}$.

ATR-mode FT-IR spectroscopic studies were performed in order to elucidate the chemical composition and structural integrity of the formed collagen hydrogel bio-constructs, namely, (i) control hydrogel construct without any drug or nanomaterials imbibed in it, (ii) citrate stabilized GNP1 entrapped hydrogel and (iii) bioconjugated GNP1-ACV entrapped collagen hydrogel. The distinct characteristics amide bands displayed by collagen via FT-IR spectral analysis can reveal an intriguing insight into its triple helical structural integrity[7] as shown in Table IV[89]. The table clearly demonstrate that the cross-linked collagen hydrogel bio-constructs synthesized does not show any detectable band shift in comparison to the band positions of native collagen reported earlier[10]. This information leads to one of the most important observations that the in spite of the cross-linking agents and drug/nanomaterials entrapped inside the collagen hydrogel, the triple helical organisation of the collagen chains remains intact in the above mentioned bio-constructs. In addition to the unchanged peak/band positions of distinct amide bands, we have also calculated the ratio of absorbance between amide$_{III}$ to 1450 $cm^{-1}$ ($Abs_{III}/Abs_{1450}$) band. This ratio particularly depicts the degree of triple helix preservation following the formation of cross-linked collagen hydrogel constructs[9][10]. A ratio of ($Abs_{III}/Abs_{1450}$) (Table V, Figure S2) closer to unity provide evidence of the preserved triple helical integrity of the formed collagen networks, which is an integral aspects of our study because if the structural integrity of the collagen hydrogel bio-constructs is compromised following the entrapment of conjugated/unconjugated GNP1, then it will lead to an instant graft rejection.

**Table IV:** Peak position and band vibrations of different collagen hydrogel bio-constructs

| | **Hydrogel-Control** | **Hydrogel-GNP1** | **Hydrogel-GNP1-ACV** | **Chemical composition** |
|---|---|---|---|---|
| **Peak position ($cm^{-1}$)** | 3294 and 3100 | 3289 and 3103 | 3296 and 3080 | Amide A and B bands-associated with the stretching vibrations of N-H groups[4] |
| **Peak position ($cm^{-1}$)** | 1634 and 1547 | 1647 and 1549 | 1632 and 1547 | Amide I and II bands- these primarily results from the stretching vibrations of peptide C=O groups and as well as from N-H bending and C-N stretching vibrations[4] |
| **Peak position ($cm^{-1}$)** | 1238 | 1238 | 1238 | Amide III band- this is assigned to the C-N stretching vibrations as well as N-H bending vibrations from the amide linkages[4]. |
| **Peak position ($cm^{-1}$)** | 1450 | 1452 | 1452 | |

**Table V:** Ratio of absorbance at 1250 $cm^{-1}$ and 1450 $cm^{-1}$

| | **Hydrogel-Control** | **Hydrogel-GNP** | **Hydrogel-GNP-ACV** |
|---|---|---|---|
| **Ratio ($Abs_{III}/Abs_{1450}$)** | 1.026 | 1.017 | 0.966 |

**Section 4: Properties of Tx implants**

The functional integrities and implicit requisites of the formulated Tx implants were tested and compared with the control (Bs, without nanosystems). Physical and biocompatibility properties of the implants are described in Figure S8.

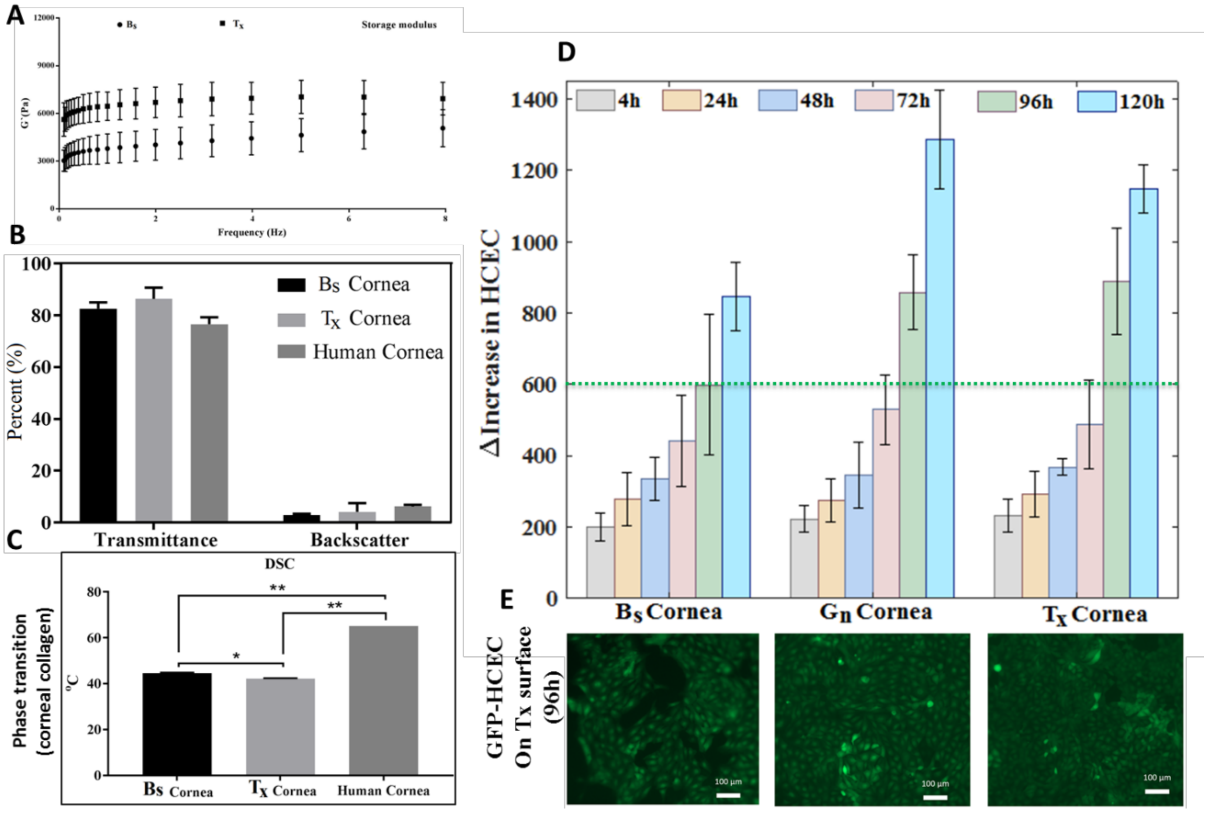


**Figure S8: Functional characterization of Tx implants compared to unmodified controls**. (**A**) Storage modulus of the samples (**B**) Optical transmittance and back scattering of control and Tx implants(**C**) Differential scanning calorimetry (DSC) of implants (**D**) Proliferation rates of human corneal epithelial cells (HCECs) on Tx, unmodified control implants (Bs) and implants with GNPs only without drugs (Gn cornea) (**E**) Fluorescence microscopic images of GFP-HCECs layer on the different implants at 96 hours post-seeding.

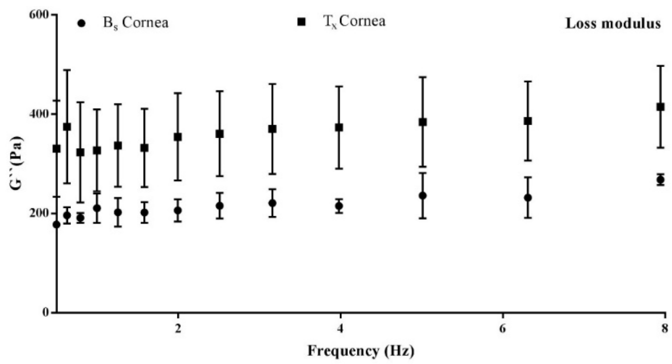


Viscoelastic properties of the hydrogels were tested with rheometer. The storage modulus (G′) is the ability to store deformation energy in an elastic manner during shear process of material.

**Figure S9.** Comparative Loss modulus

This is directly proportionate to the degree of cross-linking, meaning that the higher the degree of cross-linking correlates with the greater storage modulus. In other way, higher G' denotes more solid-like property, higher strength and mechanical rigidity. Comparing control (Bx) to Tx showed that, Tx had high storage modulus proving that Tx was mechanically strong than Bx, although denaturation temperature of the Bx was high then Tx.

#### Section 5: Growth promotion activity of GNPs on human corneal epithelial cells

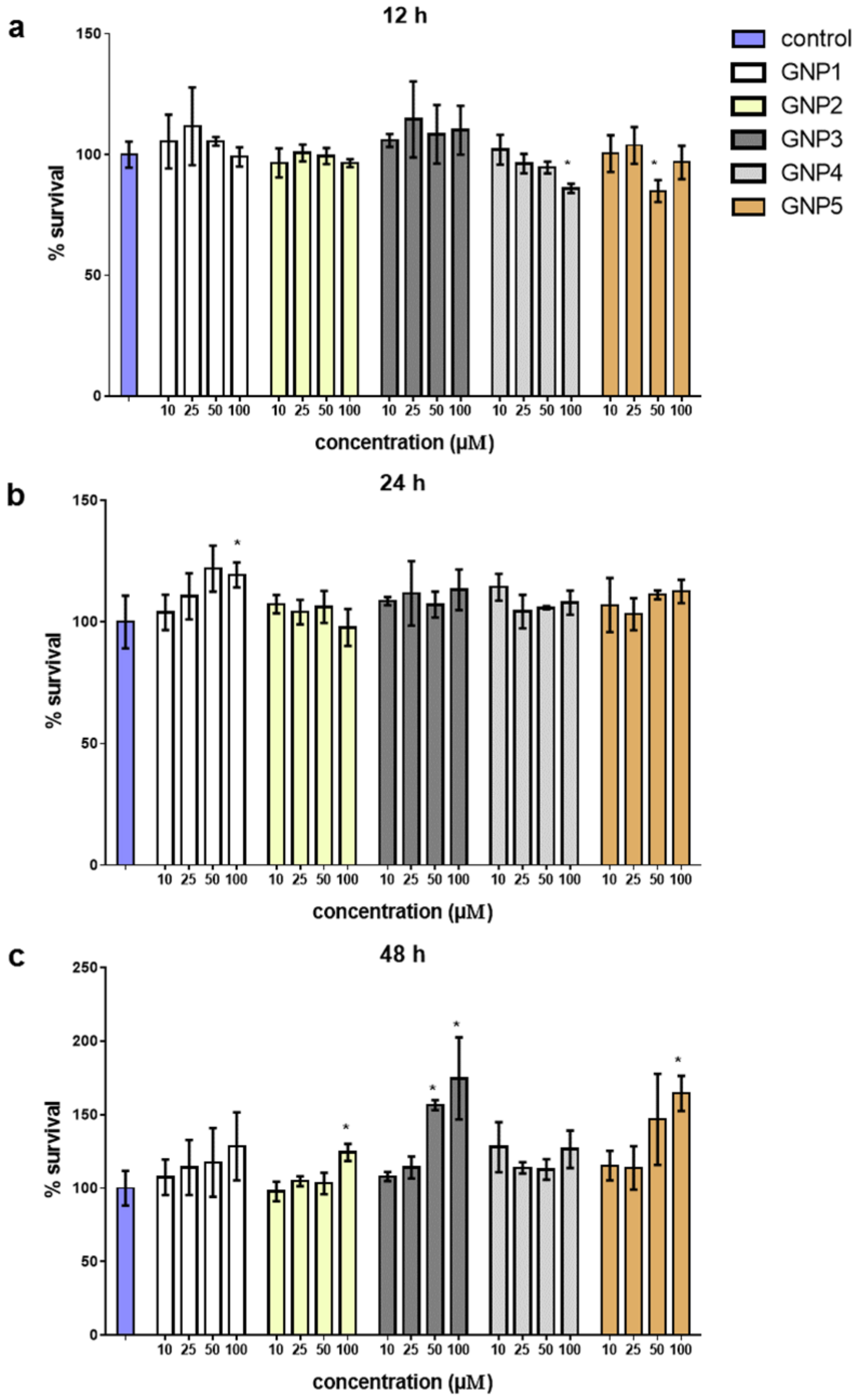


MTS assay was performed to estimate the growth promoting rate of HCECs in the presence of different concentrations of the GNPs (10, 25, 50 and 100 μM). Regardless the incubation period (12, 24 or 48 h), cell survival followed a similar pattern. No significant cytotoxicity observed with any of the GNPs examined (Figure S8 a-c).

**Figure S10.** GNP Proliferation. Percentage survival of HCECs after incubation with increasing concentrations of the different GNPs for (a) 12 h (b) 24 h and (c) 48 h. Cell viability was assessed by the MTS assay and the absorbance was recorded at 490 nm. Data are presented as mean ± standard deviation (n = 3). Unpaired t-test was performed and significance was defined as $p < 0.05$. The asterisk * indicates significant difference compared with the control. Remarkably, GNPs 2, 3 and 5 demonstrated a higher percentage of cell survival for the highest concentration.

**Section 6: Biocompatibility of FeNPs on HCECs**

Cell viability (MTS) assay was performed on HCECs in the presence of increasing concentrations of the FeNPs (10, 25, 50 and 100µM) for 12, 24 and 48 hrs. In spite of the increasing dose of FeNPs, cell survival percentage for HCECs remains the same, with more than 80% survivality even after 48hrs of incubation. No observable cytotoxity was observed for the nanoparticles (Figure S11). This result indicates that FeNPs can be a reasonable candidate for theranostic applications.

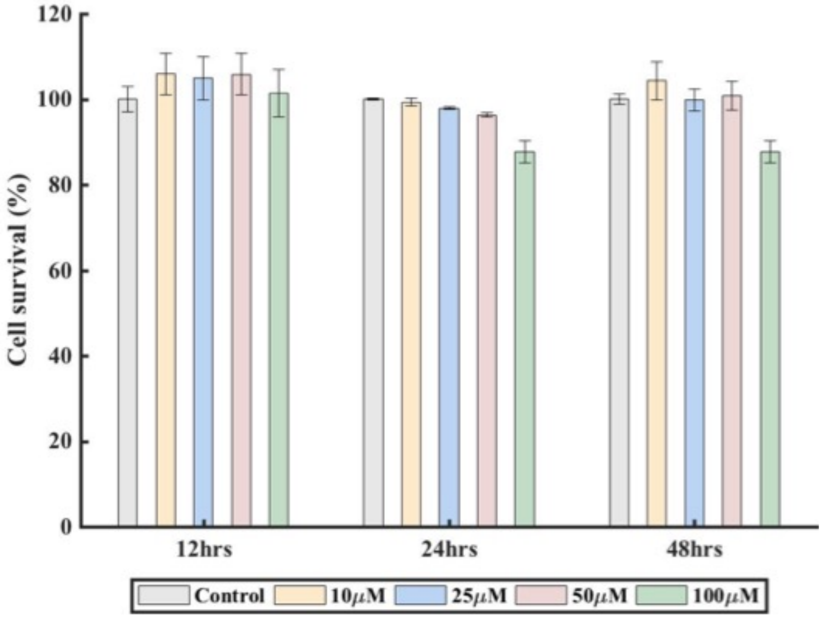


**Figure S11.** FeNPs cytotoxicity assay: Percentage survival of HCECs after incubation with increasing concentration of FeNPs from 10-100µM for 12, 24 and 48 h. MTS assay were performed for cell viability assay and absorbance was recorded at 490nm. We have optimized the safe concentration of FeNPs for the design of theranostic biosynthetic corneas. Data are presented as mean ± standard deviation (n=3). Unpaired t-test was performed and significance was defined as $p<0.05$.

#### Section 7: Potential and prospective applications of Tx implants in Corneal pathogenesis

By 2019, it is estimated that 84% of all visual impairment will be among those aged 50 years or more.

| Diseases | Broad reason | Specific reason | Treatment option | Potential Tx remedy | Tried in this paper |
|---|---|---|---|---|---|
| Infections | | | | | |
| | Bacterial | *Pseudomonas aeruginosa, Viridans streptococci, Staphylococcus aureus* | Anti-bacterial/ Artificial cornea transplantation | Yes | |
| | Fungal | *Candida albicans, Fusarium solani* | Anti-fungal/ Artificial cornea transplantation | Yes | |
| | Viral | Herpes simplex virus (HSV) | Anti-viral/ Artificial cornea transplantation | Yes | Yes |
| Trauma | | | | | |
| | Exposure to toxic chemicals | Alkali burn, acid burn | Artificial cornea transplantation | Yes | |
| Dystrophies and degenerative corneal disorders | | | | | |
| | Fuchs' dystrophy | Dysfunctional endothelium | Artificial cornea transplantation | Yes | |
| | map-dot-fingerprint dystrophy | Developmental defect of epithelium's basement membrane | Artificial cornea transplantation | Yes | |
| Ectasia (thinning) | | | | | |
| | Keratoconus | Normally round cornea becomes thin and develops a cone-like bulge | Artificial cornea transplantation | Yes | |
| Stevens-Johnson syndrome | | | | | |
| | Toxic epidermal necrolysis | Immune-complex–mediated hypersensitivity complex | Early recognition and withdrawal of all potential causative drugs | Yes/No | MRI monitoring could be useful |
| Human donor cornea transplantation associated diseases | | | | | |
| | Disease transmission | Transmission of HSV from donor to host | Artificial cornea transplantation | Yes | |
| | Immunosuppression | Compromise immune system, Activate latent HSV | Artificial cornea transplantation | Yes | |
| | Reactivation of pathogen | Reactivation of HSV during transplantation | Transplant containing anti-microbial agent | Yes | Yes |
| Treatment approach related complications | | | | | |
| | Requirement of huge dose of drug | Less amount of drug reach at the point of interest | Nano-carrier based drug delivery | Yes | Yes |
| | High dosage regimen | To keep effective concentration | Sustained release drug delivery | Yes | Yes |
| | Systemic side effect | Gastrointestinal upset | Devilry of drug from carrier through artificial cornea | Yes | Yes |
| | Resistance/latency formation by pathogen | Inefficient amount of drug at the site of action | Local drug delivery | Yes | Yes |
